# CARD-like domains mediate anti-phage defense in bacterial gasdermin systems

**DOI:** 10.1101/2023.05.28.542683

**Authors:** Tanita Wein, Alex G. Johnson, Adi Millman, Katharina Lange, Erez Yirmiya, Romi Hadary, Jeremy Garb, Felix Steinruecke, Aidan B. Hill, Philip J. Kranzusch, Rotem Sorek

**Author notes:** These authors contributed equally to this study (Tanita Wein, Alex G. Johnson).

## Abstract

Caspase recruitment domains (CARDs) and pyrin domains are important facilitators of inflammasome activity and pyroptosis. Upon pathogen recognition by NLR proteins, CARDs recruit and activate caspases, which, in turn, activate gasdermin pore forming proteins to and induce pyroptotic cell death. Here we show that CARD-like domains are present in defense systems that protect bacteria against phage. The bacterial CARD is essential for protease-mediated activation of certain bacterial gasdermins, which promote cell death once phage infection is recognized. We further show that multiple anti-phage defense systems utilize CARD-like domains to activate a variety of cell death effectors. We find that these systems are triggered by a conserved immune evasion protein that phages use to overcome the bacterial defense system RexAB, demonstrating that phage proteins inhibiting one defense system can activate another. We also detect a phage protein with a predicted CARD-like structure that can inhibit the CARD-containing bacterial gasdermin system. Our results suggest that CARD domains represent an ancient component of innate immune systems conserved from bacteria to humans, and that CARD-dependent activation of gasdermins is conserved in organisms across the tree of life.

## Introduction

Pyroptosis is a cell-autonomous process where pore-forming proteins named gasdermins control cell death in response to infection^1, 2^. In animals, this process is initiated once a pathogen is detected intracellularly by a pathogen receptor of the nucleotide oligomerization domain (NOD)-like receptor (NLR) protein family^3^. Once the NLR detects a signature of pathogen invasion via its C-terminal repeats domain, it oligomerizes to initiate the formation of an inflammasome complex^4, 5^. This complex recruits and activates proteases of the caspase family^6^, via protein–protein interactions between a caspase recruitment domain (CARD) at the N-terminus of the caspase, and a CARD domain in a caspase-recruiting adaptor protein called ASC that also interacts with the NLR^5^. Activated caspases then proteolytically cleave off a C-terminal repressor domain from a protein called gasdermin, liberating the gasdermin N-terminal domain to assemble into large pores in the cell membrane^7, 8^. These pores release pro-inflammatory cytokines and promote death of the infected cell^9^.

It was recently shown that gasdermin proteins are also present in multiple bacteria, where they are almost always encoded in an operon together with a protease-encoding gene^10^. Similar to the domain architecture of human gasdermins, bacterial gasdermins have a C-terminal inhibitory domain that can be cleaved by the associated protease, triggering the bacterial gasdermin to oligomerize into large, membrane-breaching pores^10^. A gasdermin-containing four-gene operon from *Lysobacter enzymogenes* was shown to protect against phage infection when heterologously expressed in *E. coli*^10^, but the mechanism by which the gasdermin system defends bacteria against phage remained unknown.

By studying the *Lysobacter* gasdermin system at the single-cell level *in vivo*, we show here that in response to phage infection, the *Lysobacter* gasdermin is activated by proteolytic cleavage, forms discrete puncta, and permeabilizes the cell membrane. This process leads to cell death at a time point earlier than that necessary for the phage to complete its replication, such that no mature phages are released from the infected cells. We find that gasdermin activation and maturation depends on a CARD-like domain that is encoded at the N-terminus of the associated protease, pointing to an ancient evolutionary origin for CARD domains as mediators of pyroptosis. We detect multiple additional defense systems with CARD-like domains in which the gasdermin gene is replaced by other cell-death effectors and show that the CARD-like domain is also essential for phage defense in these systems. Some phages are shown to encode CARD-only proteins that can disrupt the activity of the gasdermin system. Finally, we find that phages can escape gasdermin defense by mutating RIIB, a protein conserved in T-even phages, and that RIIB expression is sufficient for the activation of the *Lysobacter* gasdermin system.

## Results

### Single-cell analysis of gasdermin-mediated immunity in bacteria

It was previously suggested that gasdermin-mediated defense in bacteria involves premature death of infected cells^10^. To examine this hypothesis at the single-cell level we mixed two populations of *E. coli* cells, one expressing the WT *Lysobacter* gasdermin system and the other expressing GFP instead. The mixed population was infected with phage T6 and observed by time-lapse microscopy. The GFP-encoding negative control cells became lysed by the phage after ∼100 min from the onset of infection (Figure 1a). However, cells encoding the *Lysobacter* gasdermin system became membrane impaired starting 45 minutes after the onset of infection, as observed by the penetration of propidium iodide into the cells (Figure 1a; Movie S1). These cells remained membrane impaired throughout the experiment and did not grow, suggesting that gasdermin-mediated membrane permeability caused cell death prior to the completion of the phage replication cycle. In agreement with this observation, phages did not replicate when infecting gasdermin-containing cells (Figure 1b). These results establish the gasdermin system as a bacterial abortive infection system^11^ that prevents the propagation of phage and its spread to nearby cells by causing regulated death of the infected cell.

**Figure 1.**
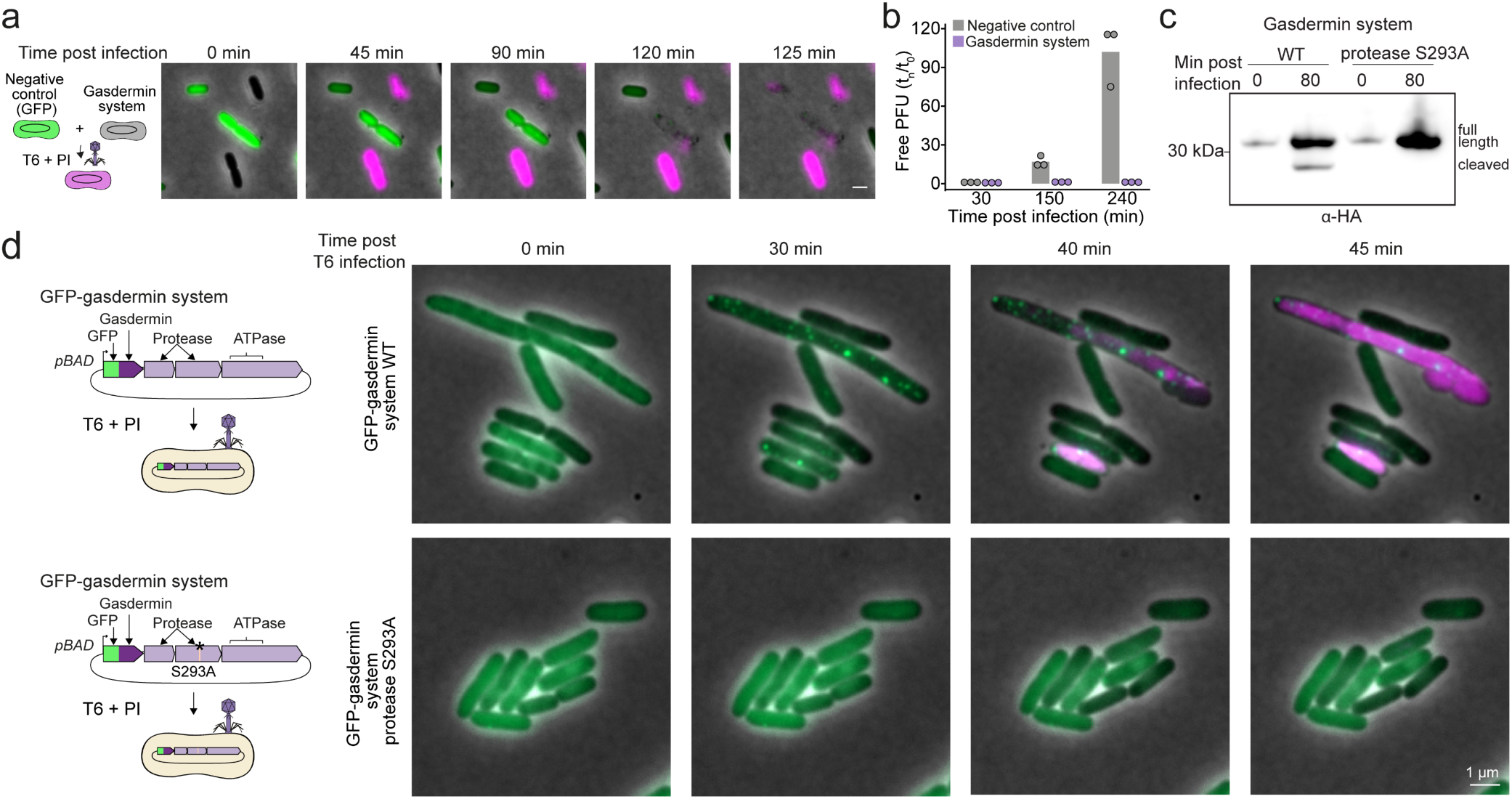
Single cell analysis of gasdermin-mediated defense in bacteria. **a.** Time-lapse microscopy of live *E. coli* cells expressing GFP (green) mixed with cells expressing the *Lysobacter* gasdermin system (black). Cells were infected with phage T6 at an MOI of 2 in the presence of propidium iodide (PI) and incubated at room temperature on an agar pad. Overlay images of phase contrast, green channel (GFP), and magenta channel (PI) are presented (scale bar = 0.5 µm). **b.** Phage replication assay. Plaque-forming units of phage T6 were sampled from the supernatant of *E. coli* cells containing an empty vector as a negative control or expressing the gasdermin system. Data show measured phage titer divided by the titer measured prior to infection. Cells were infected at an MOI of 0.01. Bars represent the average of three replicates with individual data points overlaid. **c.** Gasdermin cleavage during phage infection. Western blot analysis of N-terminally HA-tagged gasdermin following infection by phage T6 at an MOI of 2, in the WT or protease-mutated *Lysobacter* gasdermin system. **d**. Time-lapse microscopy of live cells expressing the WT or mutated *Lysobacter* gasdermin system in which gasdermin was N-terminally fused to GFP. Cells were infected with phage T6 in the presence of propidium iodide (PI) and incubated at room temperature on an agar pad. Overlay images of phase contrast, green channel (GFP), and magenta channel (PI) are presented.

The proteases associated with bacterial gasdermins were shown *in vitro* to cleave the bacterial gasdermin, thus activating it to become a pore-forming protein^10^. Our data now show that gasdermin proteolytic cleavage is specifically induced by phage infection, as demonstrated by Western blot analysis of proteins extracted from infected cells (Figure 1c, Figure S1). To further examine whether activation of gasdermins leads them to aggregate *in vivo*, we visualized *E. coli* cells encoding the *Lysobacter* gasdermin system, in which the gasdermin protein was N-terminally tagged with GFP. Prior to phage infection gasdermins appeared evenly spread within these cells; but following infection gasdermin proteins formed multiple clear puncta suggesting that they were activated to form membrane pores (Figure 1d; Movie S2). Indeed, cells in which gasdermin puncta were observed became membrane permeated, as inferred by propidium iodide staining (Figure 1d; Movie S2). Combined, these results show that similar to the activity of their human counterparts, bacterial gasdermins are activated by proteolytic cleavage in response to infection and then aggregate to form membrane pores that cause cell death.

### CARD-like domains are essential for bacterial gasdermin-mediated defense

To gain further insight into the regulation of gasdermin activation in bacteria we examined the genes in the *Lysobacter* gasdermin operon. This operon comprises, in addition to the gasdermin gene, two consecutive genes encoding proteins with trypsin-like protease domains, and a gene encoding a protein annotated as an ATPase (Figure 2a). It was previously shown that intact active sites of the second protease and the ATPase, but not of the first protease, are necessary for defense against phages^10^, suggesting that the first protease gene is dispensable for defense against the phages tested here. A closer examination of the domain architecture of the ATPase showed that it belongs to the STAND ATPase protein family, which are bacterial proteins that are considered the evolutionary ancestors of eukaryotic NLRs^12–14^ (Figure 2a; Figure S2). It was recently shown that bacterial STAND ATPases are pattern recognition proteins that, similar to human NLRs, detect infection via their C-terminal repeat region and oligomerize into an active multimeric complex once they bind their ligand^14^. The *Lysobacter* gasdermin operon architecture therefore implies that pyroptosis in bacteria may be initiated by a pattern recognition NLR-like protein, in similarity to pyroptosis regulation in human cells.

**Figure 2.**
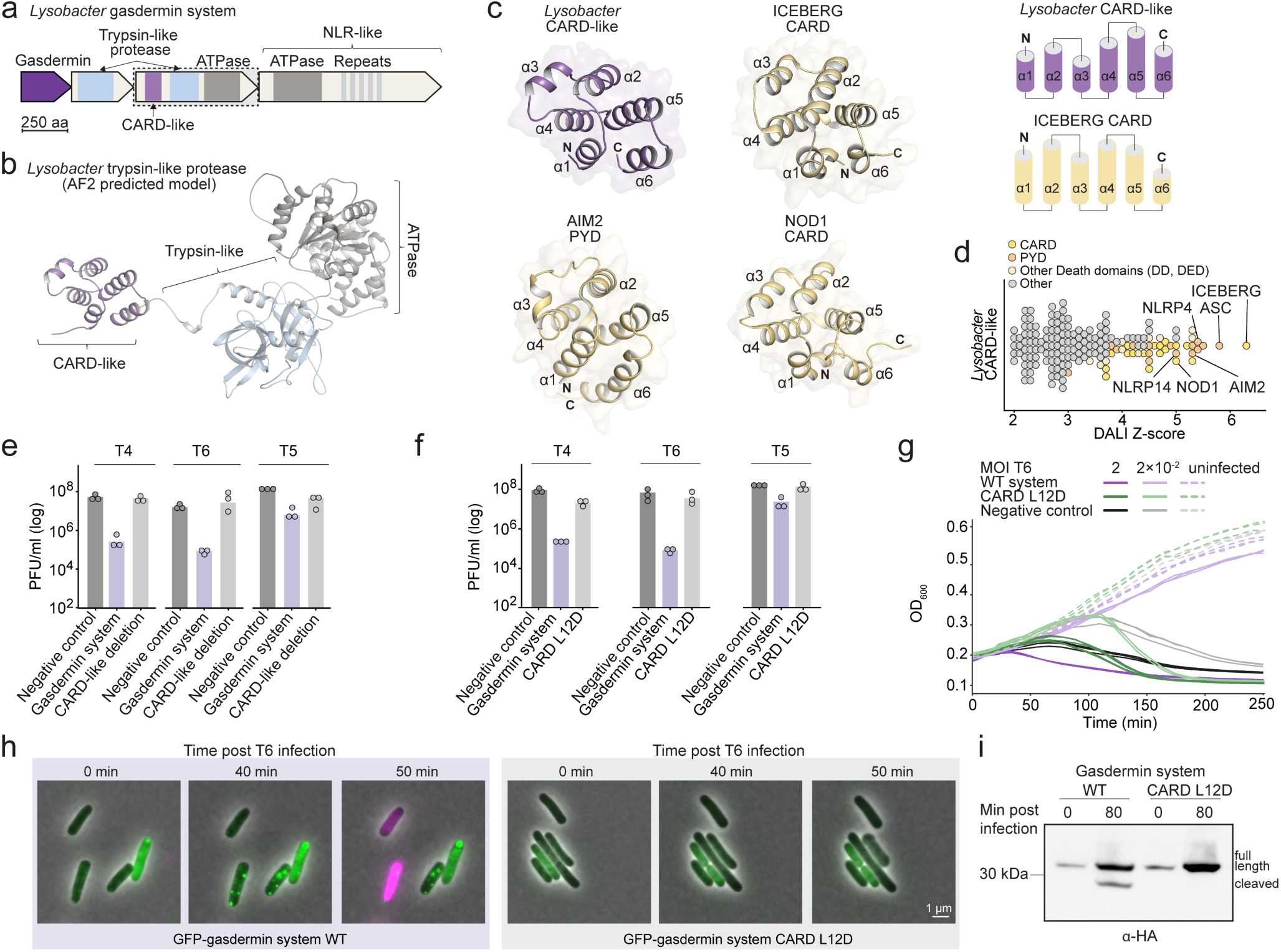
CARD-like domains in the bacterial gasdermin system. **a.** Domain architecture of the gasdermin defense system from *Lysobacter enzymogenes*. **b.** Alphafold2 prediction of the protease with the CARD-like domain. **c**. Crystal structure of the CARD-like domain encoded in the N-terminus of the *Lysobacter* gasdermin-associated protease. Also shown, for comparison, are the human ICEBERG CARD domain (PDB ID 1DGN), the PYD domain from human AIM2 (PDB ID 2N00) and the CARD domain from human NOD1 (PDB ID 2DBD) as well as topology diagrams of the *Lysobacter* CARD-like domain and the ICEBERG CARD domain. **d.** DALI Z scores of protein structures similar to the structure of the *Lysobacter* CARD-like domain. **e**. The CARD like domain is essential for gasdermin-mediated defense. Efficiency of plating of phages infecting *E. coli* cells that express either the WT gasdermin system or the system in which the CARD-like domain (residues 1-91) was deleted. Data represent plaque-forming units (PFU) per ml. Average of three independent replicates, with individual data points overlaid. Negative control is a strain in which GFP is expressed instead of the gasdermin system. **f.** A mutation in a conserved residue in the CARD-like domain abolishes defense. Experiment was performed as in panel E. **g**. Liquid culture growth of *E. coli* cells with WT or CARD-mutated gasdermin system, infected by phage T6 at 25°C. Bacteria were infected at time 0 at an MOI of 2 or 0.02. Three independent replicates are shown for each MOI, and each curve represents an individual replicate. **h.** Time-lapse microscopy of live *E. coli* cells expressing the WT or mutated *Lysobacter* gasdermin system in which gasdermin was N-terminally fused to GFP. Cells were infected with phage T6 in the presence of propidium iodide (PI) and incubated at room temperature on an agar pad. Overlay images of phase contrast, green channel (GFP), and magenta channel (PI) are presented. **i.** Gasdermin cleavage depends on integrity of a CARD-like domain. Western blot analyses of N-terminally HA-tagged gasdermin following infection by phage T6 at an MOI of 2, in the WT or CARD-mutated *Lysobacter* gasdermin system.

While analyzing the domain architecture of the essential protease in the system we noticed, in addition to the protease domain, a non-annotated domain of approximately 100 amino acids at the extreme N-terminus (Figure 2a). Analysis of this domain using AlphaFold^15^ suggested the presence of an α-helical region (Figure 2b) that we hypothesized may have similarity to human α-helical CARD domains essential for inflammasome signaling. To test a possible evolutionary connection with human immunity, we expressed the N-terminus of the gasdermin-associated protease in isolation and determined a 1.25 Å crystal structure of the protein domain (Figure 2c,d; Table S1). The structure revealed striking homology with human CARD and PYD domains including shared conservation of each of the six α-helices common to members of the Death domain protein superfamily^16^.

CARD and PYD domains are essential for inflammasome activity and apoptosis in humans, participating in the assembly of large immune protein complexes via homotypic protein-protein interactions^17, 18^. These domains are usually present in the N-termini of pattern recognition immune receptors such as NLRs, RIG-I and AIM2; in immune adaptor proteins such as ASC and MyD88; and in the N-termini of caspases involved in immune-related cell death^16, 19^, similar to the positioning of the bacterial CARD domain in the *Lysobacter* protease (Figure 2b). Using DALI structural comparison^20^, we observed that the structure of the *Lysobacter* CARD-like domain is most similar to the CARD domain in the human ICEBERG protein^21^ as well as PYDs of human ASC, NLRP4, and AIM2 proteins (Figure 2c,d). Notably, the *Lysobacter* CARD-like domain shares identical topology with the human proteins ICEBERG, AIM2, and NOD1 with each of the 6 α-helices exhibiting near identical angular positioning and packing within the alpha-helical bundle (Figure 2c,d).

Deletion of the CARD-like domain from the protease gene resulted in loss of anti-phage defense, suggesting that this domain is essential in the gasdermin defense system (Figure 2e). We next designed a point mutation within the bacterial CARD-like domain based on structural comparisons with human CARD and PYD domains (Figure S3). A mutation in residue L10 of the PYD domain from the human AIM2 inflammasome was previously shown to disrupt PYD self-association required for inflammasome assembly^22^. A mutation in the structurally equivalent L12 residue in the *Lysobacter* CARD-like domain abolished anti-phage defense (Figure 2f).

Infection of bacteria in liquid culture showed that cells encoding the gasdermin system with the L12D mutation in the CARD-like domain were not protected and did not undergo premature cell death (Figure 2g). Microscopy experiments confirmed that these cells do not become membrane impaired in response to infection (Figure 2h). The gasdermin protein in these cells remained evenly spread in the cytoplasm and did not form puncta, suggesting that the CARD-like domain is essential for the maturation of the gasdermin protein (Figure 2h). Indeed, the gasdermin protein was not cleaved by the protease when the N-terminal CARD of the protease was mutated (Figure 2i). These results show that the CARD-like domain is essential for gasdermin cleavage and maturation by the associated protease during anti-phage defense in bacteria.

### Multiple anti-phage defense systems encode CARD domains

Our discovery of a bacterial defense mechanism that depends on a CARD-like domain prompted us to investigate whether there may be additional phage resistance systems that utilize CARD-like domains. Sequence-based homology searches identified multiple bacterial proteins with similarity to the *Lysobacter* CARD-encoding protease (Table S2). In all cases, these proteases were encoded in operons similar to the *Lysobacter* gasdermin system, comprising two proteases as well as the NLR- like protein (Figure 3a). In most cases, AlphaFold2 analysis of the domains in the N-termini of the second protease suggested clear predicted homology to the experimental crystal structure of the *Lysobacter* CARD-like domain (Figure S4).

**Figure 3.**
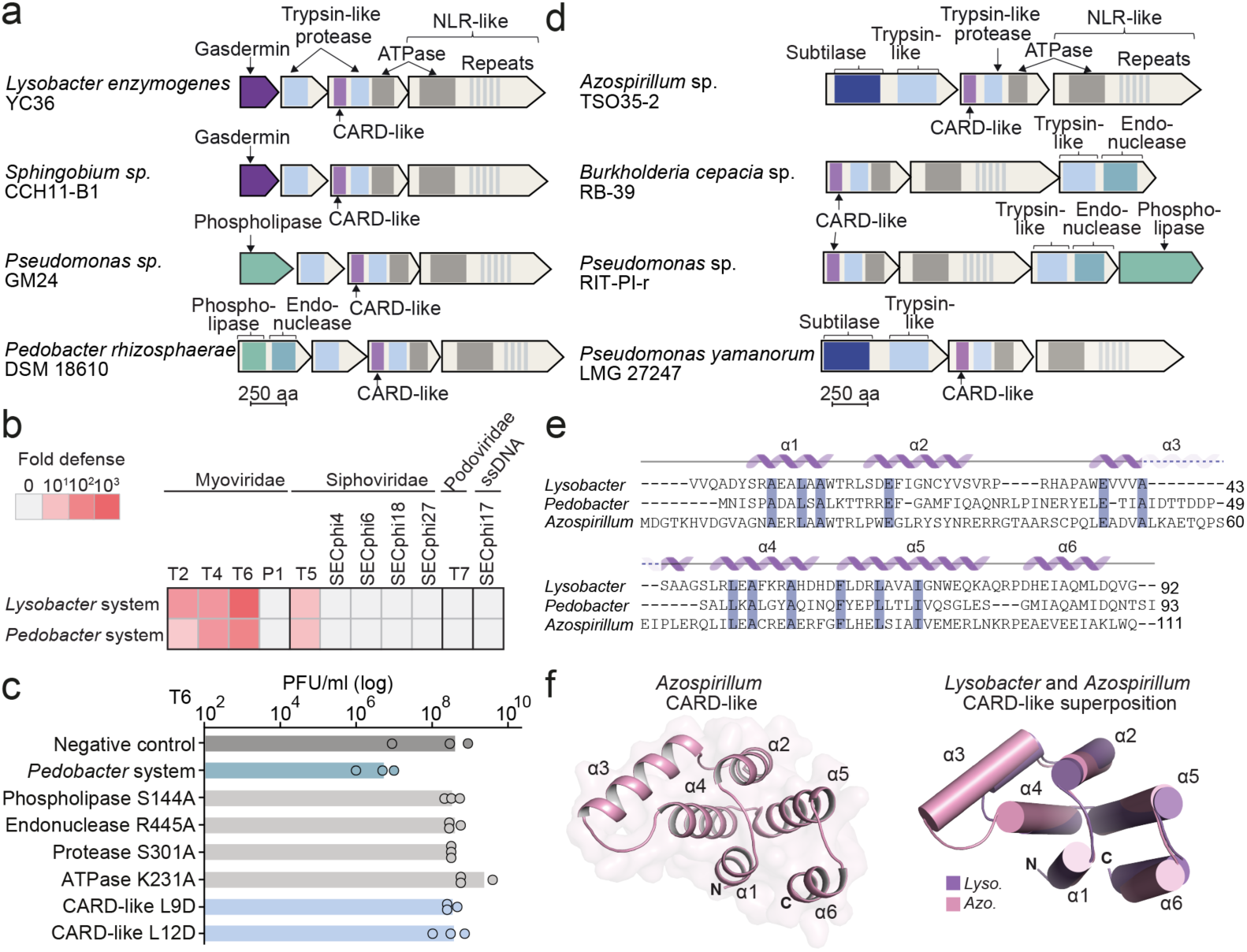
CARD-like domains in multiple bacterial defense systems. **a**. Domain architecture of operons homologous to the *Lysobacter* gasdermin system. **b.** Defense profiles of the *Lysobacter* and the *Pedobacter* operons transformed into *E. coli* under an inducible promoter. Fold defense was measured using serial dilution plaque assays, comparing the efficiency of plating (EOP) of phages on the system-containing strain with the EOP on a control strain that lacks the system. Data represent an average of three replicates (see Figure S5a). **c.** EOP of phages infecting *E. coli* cells that express either the WT or mutated system from *Pedobacter rhizosphaerae*. Data represent PFU per ml. Average of three independent replicates, with individual data points overlaid. Negative control is a strain in which GFP is expressed instead of the defense system. **d.** Domain architecture of operons in which structural homologs of the *Lysobacter* and *Pedobacter* proteases were detected. **e.** Multiple sequence alignment of the CARD-like domain from the protease of the *Lysobacter*, *Pedobacter* and *Azospirillum* systems. Shading indicates residue conservation and secondary structure elements are based on the *Lysobacter* CARD-like domain. **f.** Crystal structure of the CARD-like domain encoded in the N-terminus of the *Azospirillum* protease, and superposition of the *Azospirillum* and *Lysobacter* CARD-like domains.

Intriguingly, in most of these systems the gasdermin-encoding gene was replaced by another gene (Figure 3a). These gasdermin-replacing genes encoded proteins with endonuclease and phospholipase domains, both of which often function as effectors in other bacterial defense systems that protect via abortive infection^23^ (Figure 3a). We selected such a system from *Pedobacter rhizosphaerae*, encoding an effector protein with an N-terminal phospholipase domain and a C-terminal endonuclease domain, for further experimental examination. This system conferred defense against the same set of phages as the *Lysobacter* gasdermin system, implying that it may sense the same signature of phage infection despite encoding a different effector (Figure 3b, Figure S5a). Similar to the *Lysobacter* gasdermin system both the ATPase gene as well as the protease were essential for anti-phage defense (Figure 3c). Point mutations in either the phospholipase or endonuclease domains of the effector abolished defense, suggesting that both domains are involved in the effector function (Figure 3c). Point mutations in the conserved residues in the CARD-like domain of the *Pedobacter* system showed that this domain is also essential for defense in this system (Figure 3c).

To extend the search for CARD-encoding defense systems, we used FoldSeek^24^ to detect distantly homologous proteins that share a similar fold as the CARD-containing proteases of the *Lysobacter* and *Pedobacter* defense systems. This search revealed a family of three-gene operons that encode the CARD-containing protease, an ATPase with C-terminal repeats, and predicted effector proteins that are often fused to trypsin-like proteases (Figure 3d; Table S3). These operons were enriched next to known defense systems, suggesting a function in defense against phage (Figure S5b). Sequence alignment between the N-termini of the proteases in these systems and the CARD-like domains from the *Lysobacter* and *Pedobacter* systems showed poor sequence similarity (Figure 3e). However, by determining a crystal structure of the N-terminal domain from one of these systems encoded in *Azospirillum sp*. TSO35-2, we found that the protein adopts a clear Death-domain fold with near identical structural similarity to the CARD-like domain of the *Lysobacter* gasdermin system, as well as to CARD and PYD domains of human immune proteins (Figure 3f; Figure S5c).

In human pyroptosis, regulation of caspase activation is mediated by CARD and PYD domains, which are encoded at the N-termini of both the pattern-recognizing NLR and the caspase. We were not able to detect a PYD or CARD domain at the N-terminus of the NLR-like gene in the *Lysobacter* gasdermin system. Rather, the N-terminus of the NLR-like protein contains a predicted alpha-helical domain of ∼150aa with no similarity to known domains (Figure S6a). This N-terminal domain was conserved in homologs of the NLR-like gene from the defense systems described above.

Although we could not find a PYD or CARD-like domain at the N-terminus of the gasdermin-associated NLR-like protein, examination of the AlphaFold2 prediction for the structure of its C-terminus surprisingly revealed a predicted CARD-like domain (Figure S6a-c). In humans, CARD domains are rarely present in the C-termini of NLR proteins, where they can serve for recruiting caspases^16, 25^. Deletion of the CARD-like domain from the C-terminus of the NLR-like protein in the *Lysobacter* gasdermin system rendered the system toxic to the encoding cells (Figure S6d), suggesting that the CARD-like domain in the bacterial NLR-like protein may be responsible for auto-repression, as was shown in the case of the PYD domain in human NLRP1^26^. Indeed, further mutating the CARD-like domain of the protease in the system reverted the toxicity induced by the deletion of the bacterial NLR-like CARD domain (Figure S6d) implying that this toxicity results from the activation of the gasdermin system.

### A conserved phage protein activates the bacterial gasdermin system

Despite the efficient protection conferred by the gasdermin defense system, we noticed that when infecting system-encoding bacteria with a high concentration of phages, a small number of plaques still formed. We hypothesized that these represent phages that escaped defense due to spontaneous mutations in the gene that is recognized by the gasdermin system as a signature for infection, as previously reported for other defense systems^27^. To test this hypothesis, we collected six T6 escaper phages and sequenced their genomes (Figure 4a). In each of the six escaper phages we detected only a single mutation as compared to the genome of the WT phage (Figure 4b). In all cases, the detected mutation was in the *rIIB* gene of the T6 phage. These mutations were either early frameshift mutations (in five escapers) or a premature stop codon (one escaper), all in *rIIB*, suggesting that eliminating the expression of this phage protein was sufficient for escape from gasdermin-mediated defense (Figure 4a,4b).

**Figure 4.**
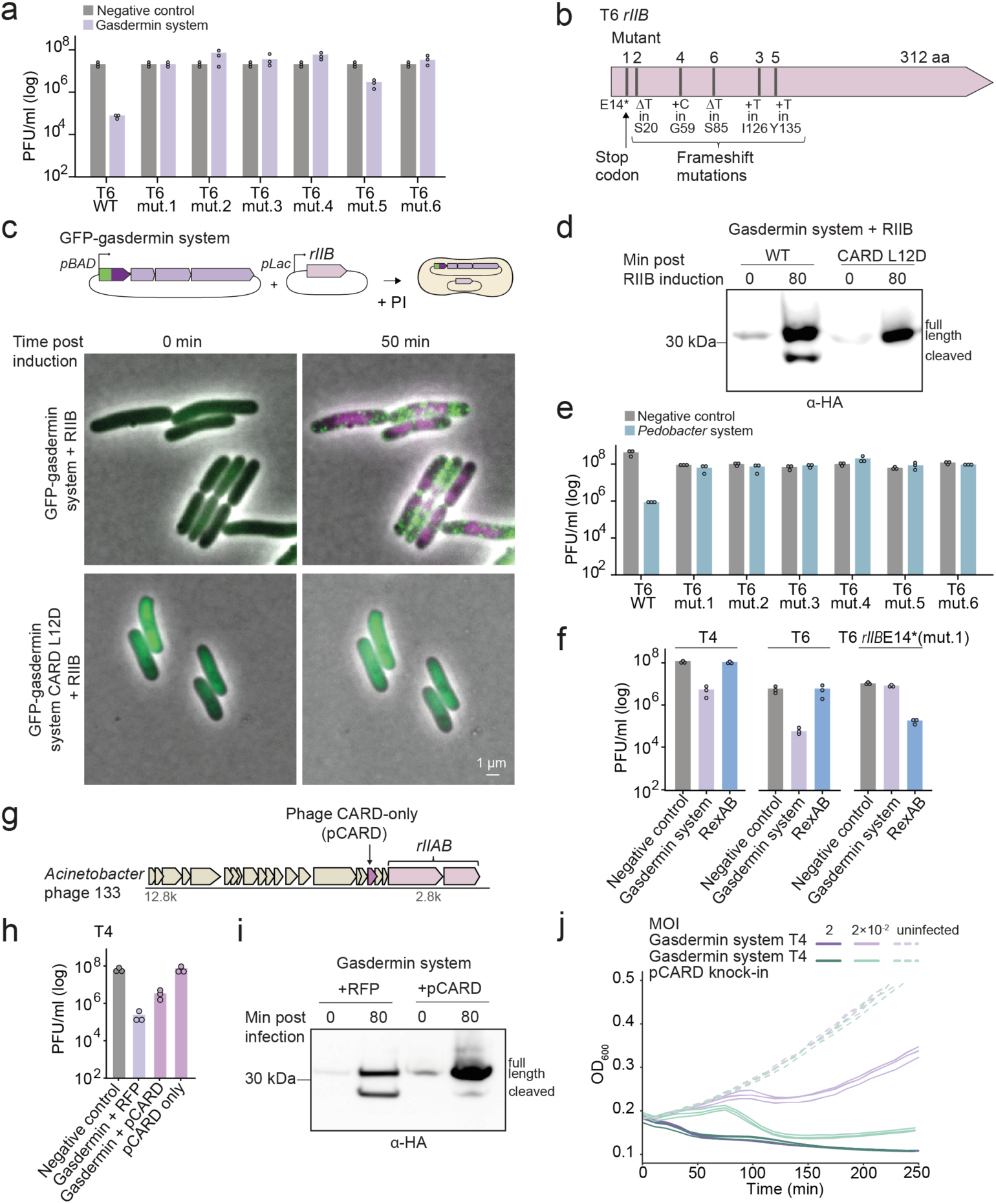
A phage protein that activates gasdermin-mediated defense. **a.** T6 mutants that escape gasdermin-mediated defense. Data represent PFU per ml of phages infecting cells expressing the *Lysobacter* gasdermin system or control cells expressing GFP instead. **b.** Positions mutated in the *rIIB* gene of phage T6 escaper phages. Mutant numbers correspond to the numbers in panel A. **c**. Phage RIIB activates gasdermin aggregation in the absence of phage infection. Time-lapse microscopy of live cells co-expressing RIIB with the WT or mutated *Lysobacter* gasdermin system, in which gasdermin was N-terminally fused to GFP. Cells were visualized at room temperature on an agar pad in the presence of propidium iodide (PI). Overlay images of phase contrast, green channel (GFP), and magenta channel (PI) are presented. **d.** Gasdermins are cleaved when the system is co-expressed with RIIB. Western blot analyses of N-terminally HA-tagged gasdermin following induction of RIIB expression, in the WT or CARD-mutated *Lysobacter* gasdermin system. **e.** EOP of T6 mutant phages infecting cells that express the *Pedobacter* defense system. **f.** EOP of phages infecting *E. coli* cells that express either the *Lysobacter* gasdermin system or the RexAB system from the phage Lambda. Data represent PFU per ml. Average of three independent replicates, with individual data points overlaid. Negative control is a strain in which GFP is expressed instead of the defense system. **g.** Genomic neighborhoods of a phage CARD-only (pCARD) protein. **h.** The phage CARD-only protein interferes with gasdermin-mediated defense. Data represent PFU per milliliter of phages infecting cells expressing the *Lysobacter* gasdermin system together with a plasmid expressing pCARD or RFP, as well as cells expressing the pCARD gene alone. Control cells express GFP. **i.** Gasdermin cleavage is reduced in the presence of pCARD. Western blot analyses of N-terminally tagged gasdermin expressed together with pCARD or RFP following infection with phage T6 at an MOI of 2. **j.** Liquid culture growth of gasdermin-expressing *E. coli* cells infected with WT T4 or with T4 engineered to encode the pCARD gene instead of its *IPI* gene. Bacteria were infected at time 0 at an MOI of 2 or 0.02 at 25 °C. Three independent replicates are shown for each MOI, and each curve represents an individual replicate.

To test if the phage *rIIB* gene alone is sufficient for triggering the gasdermin system, we cloned this gene from phage T6 under an inducible promoter and introduced it to cells containing the *Lysobacter* system in which GFP was N-terminally fused to the gasdermin protein (Figure 4c). Since RIIB expression was toxic to *E. coli* regardless of the presence of the gasdermin system (Figure S7a), we kept the cells under conditions repressing RIIB expression (1% glucose) before expressing it together with the gasdermin system. Co-expression of RIIB with the gasdermin system, in the absence of phage infection, rapidly caused gasdermin cleavage, aggregation and membrane permeabilization (Figure 4c, 4d; Movie S3). Notably, expression of RIIB together with the system mutated in the CARD-like domain of the protease did not result in gasdermin cleavage and the formation of GFP-gasdermin foci (Figure 4c, 4d). These results suggest that expression of the phage RIIB protein triggers the *Lysobacter* defense system to process gasdermin into the active pore-forming protein.

Due to the sequence homology between the ATPase and protease proteins of the *Lysobacter* and the *Pedobacter* defense systems (Figure 3a), and the fact both systems defend against the same set of phages (Figure 3b), we hypothesized that RIIB would activate the *Pedobacter* system as well. In support of this hypothesis, T6 mutants that escaped the *Lysobacter* gasdermin system were also able to infect cells expressing the defense system from *Pedobacter* (Figure 4e). Attempts to transform the *rIIB*-encoding plasmid into cells that contain the *Pedobacter* system failed, possibly indicating that expression leakage from the inducible promoters of both *rIIB* and the *Pedobacter* system resulted in activation of the system leading to cell death (Figure S7b).

The phage protein RIIB is commonly present in T-even phages, where it is encoded within an operon together with the gene *rIIA*^28^. It was previously shown that the *rIIAB* operon confers T-even phages such as T4 and T6 with the ability to overcome a defense system termed RexAB in *E. coli*^29^. T4 phage mutated in *rIIAB* cannot infect RexAB-encoding cells, while phages with intact *rIIAB* operons overcome this defense system^30^, although the mechanism by which RIIAB inhibits RexAB is unclear^31^. Our data suggest that the gasdermin defense system converged to sense the effect of the phage RIIB protein that is necessary for inhibition of another defense system (RexAB). Indeed, T4 and T6 phages that are sensitive to the gasdermin system overcame RexAB in plaque assay experiments, while mutant T6 phages that escaped the gasdermin system were sensitive to RexAB (Figure 4f). These results highlight the intricate arms race between bacteria and phages, and are reminiscent of mechanisms of defense systems such as PARIS, which becomes activated in response to phage proteins that inhibit restriction-modification systems^32, 33^.

### A phage-encoded CARD-like protein disrupts gasdermin-mediated defense

While examining the *rIIAB* locus in the T4-like *Acinetobacter* phage 133, we noticed that the *rIIAB* operon was encoded next to a small phage protein (141 aa, Figure 4g) which, based on AlphaFold2 structural analysis, was predicted to encode a CARD-like domain with no other protein domains (Figure S7c). Viruses that infect animals and humans, such as poxviruses, are known to encode pyrin-only proteins that inhibit the activation of inflammasomes by interfering with the PYD-mediated protein- protein interaction^34, 35^. As recent studies showed that phage proteins that inhibit bacterial defense systems can be encoded next to the phage protein that triggers the system^36^, we hypothesized that the phage encoded CARD-only protein may serve to interfere with the gasdermin systems.

To test this hypothesis, we co-expressed the phage CARD-only protein in cells that also expressed the gasdermin system. Plaque assay analyses showed that these cells became more sensitive to infection by phage despite encoding the gasdermin system, suggesting that the phage protein inhibits the activity of the gasdermin defense system (Figure 4h). In support of this observation, Western blot analysis of proteins from infected cells show that expression of the phage CARD-like protein results in reduced gasdermin cleavage (Figure 4i). To further examine whether this protein can confer phages with anti-defense activity, we used Cas13a^37^ to engineer the gene from *Acinetobacter* phage 133 into phage T4 under the T4 promoter naturally controlling the anti-restriction protein IPI^38^. T4 phages expressing the CARD-only protein became partially resistant to the activity of the gasdermin defense system (Figures 4j, S7d). These results suggest that the phage CARD-like protein encoded next to the *rIIAB* locus in *Acinetobacter* phage 133 inhibits the proper activity of the bacterial gasdermin system.

## Discussion

In this study we investigated the activation process of the bacterial gasdermin in response to phage infection *in vivo*, on a single-cell basis. We found that, upon infection, the bacterial gasdermin is activated by proteolytic cleavage and oligomerizes to cause premature cell death akin to mammalian gasdermins. Unexpectedly we discovered that gasdermin activation is mediated by CARD-like domains that are essential for the function of the defense system. CARD and PYD domains are known to play key roles in multiple cell death pathways, including pyroptosis, necroptosis, and apoptosis^39^. While their function has been extensively studied in mammalian innate immunity, their evolutionary origin remained elusive and was considered restricted to metazoans species^40^, although recent computational analyses suggested an earlier origin^41^. Our findings suggest that CARD domains emerged in bacteria where they originally evolved to function in antiviral defense, as recently shown for other components of the human innate immune system^42^. Our results also establish a functional connection between CARD-like domains and cell death effector proteins including gasdermins, further supporting the bacterial origin of CARD domains as mediators of cell death in response to infection.

While our results determine the presence and essentiality of CARD-like domains in the bacterial gasdermin systems, the mechanism by which these domains mediate gasdermin activation remains to be studied. In mammalian immune proteins, CARD domains and other members of the Death domain superfamily were shown to have multiple different functions. In inflammasome assembly they can recruit caspases via mediator proteins such as ASC^5^, or in some cases via direct interactions with the caspase^43^. In other cases, Death domains can be negative regulators of inflammasome activity, as in the case of the PYD domain in human NLRP1^26^. Some CARD domains were shown to be the direct sensors of pathogen infection, for example, the mouse caspase-11 CARD that directly interacts with LPS, activating caspase-11 to cleave gasdermin leading to pyroptosis^44^. Whether bacterial CARD-like domains function similar to their mammalian counterparts, or whether they have different functions in bacterial immunity, remains to be determined in future studies.

Our data show that phages can escape the bacterial gasdermin system via mutations in their *rIIb* gene, and that co-expression of RIIB alone in cells expressing the defense system is sufficient to activate gasdermin cleavage (Figure 4). These data suggest that the gasdermin system senses RIIB or its effects on the cell. RIIB was initially discovered as an anti-defense protein in T-even phages, essential for infecting bacteria that encode the defense system RexAB but dispensable for infection in cells where RexAB is absent^30^. Our findings therefore point to an intriguing arms race in bacteria, where phages encoding a protein that serves to evade one immune system become sensitive to other defense systems. Such arms race outcomes can explain why bacteria must encode multiple defense systems at the same time^45^.

In recent years, multiple components of the human innate immune system were shown to have originated from bacterial immune systems. In addition to gasdermin, these include key immune proteins such as cGAS, STING, Argonaute, TIR domains, viperin and more^42^. While these represent individual proteins that were adopted into the mammalian cell-autonomous innate immune system, our discovery of a bacterial defense system that includes gasdermin, proteases, CARD-like domains and an NLR-like protein suggests that the full inflammasome pathway leading to pyroptosis may have been originally invented in bacteria.

## Supplementary Material

### Supplementary Figures

**Figure S1.**
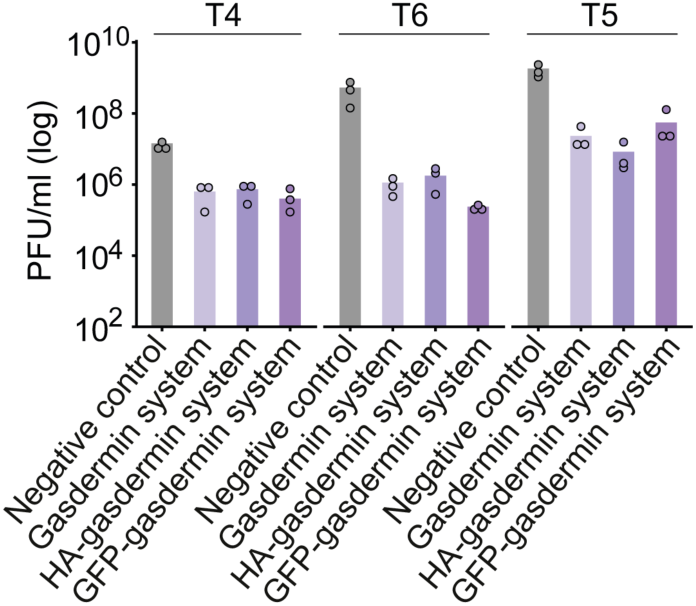
Anti-phage defense of tagged *Lysobacter* gasdermin system. Efficiency of plating of phages infecting *E. coli* cells that express the WT *Lysobacter* gasdermin system, as well as systems in which gasdermin was N-terminally fused to HA-tag or to GFP. Negative control is a strain in which GFP is expressed instead of the gasdermin system. Data represent plaque-forming units (PFU) per ml. Average of three independent replicates, with individual data points overlaid.

**Figure S2.**
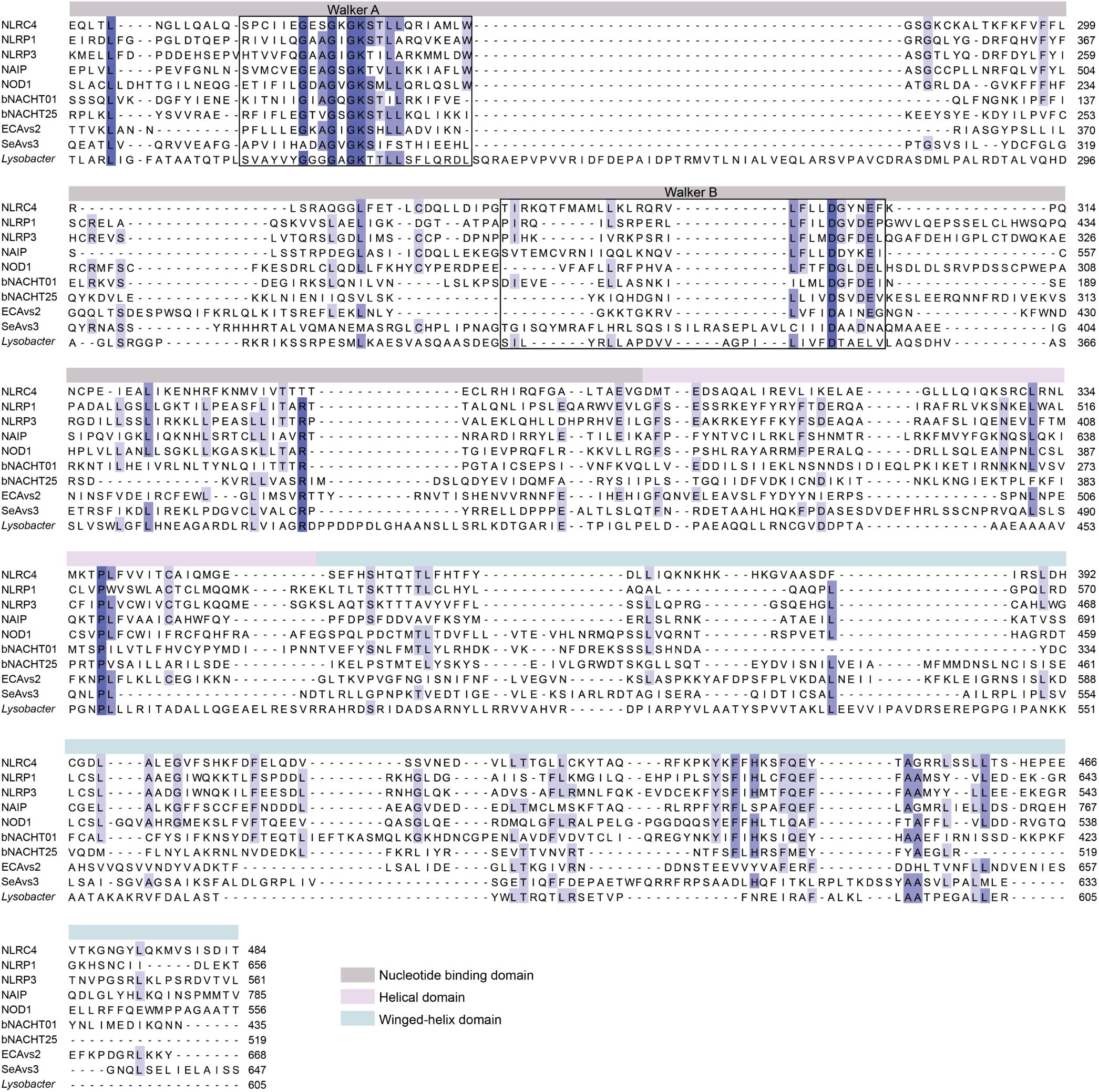
Multiple sequence alignment of the Nucleotide binding oligomerization domain (NOD) module of STAND ATPases. Shown are NOD modules from human proteins (NLRC4, NCBI accession: AAH31555; NLRP1, AAG30288; NLRP3, AAL33911; NAIP, A55478; NOD1, AAD29125) and bacterial proteins (bNACHT01, NCBI accession: WP_015632533.1; bNACHT25, WP_001702659.1; ECAvs2, WP_063118745.1; SeAvs3, WP_126523998.1), including the ATPase protein from the *Lysobacter* gasdermin system (IMG gene ID 2841794910). The depth of shading indicates degree of residue conservation.

**Figure S3.**
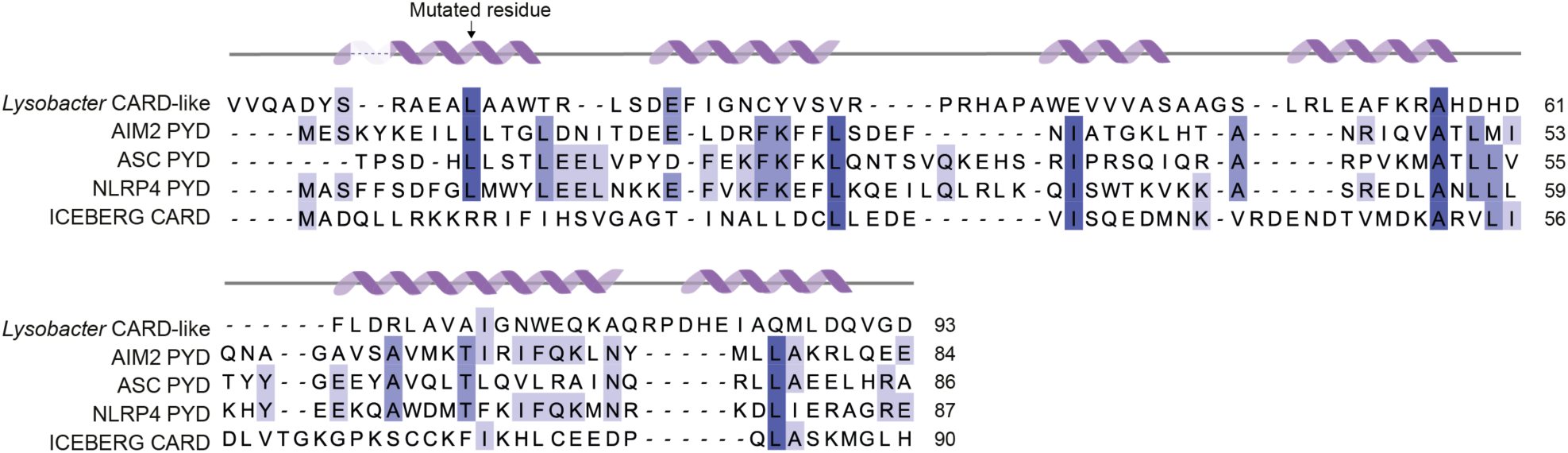
Structure-guided alignment of Death domains. Multiple sequence alignment of CARD and PYD domains from human proteins (AIM2, NCBI accession: AAB81613.1; ASC, BAA87339.2; NLRP4, AAL35293.1; and ICEBERG, AAG23528) and from the protease of the *Lysobacter* gasdermin system (gene ID 2841794909 in the IMG database^46^). The depth of shading indicates degree of residue conservation. Secondary structure elements are indicated based on the *Lysobacter* protease CARD-like structure.

**Figure S4.**
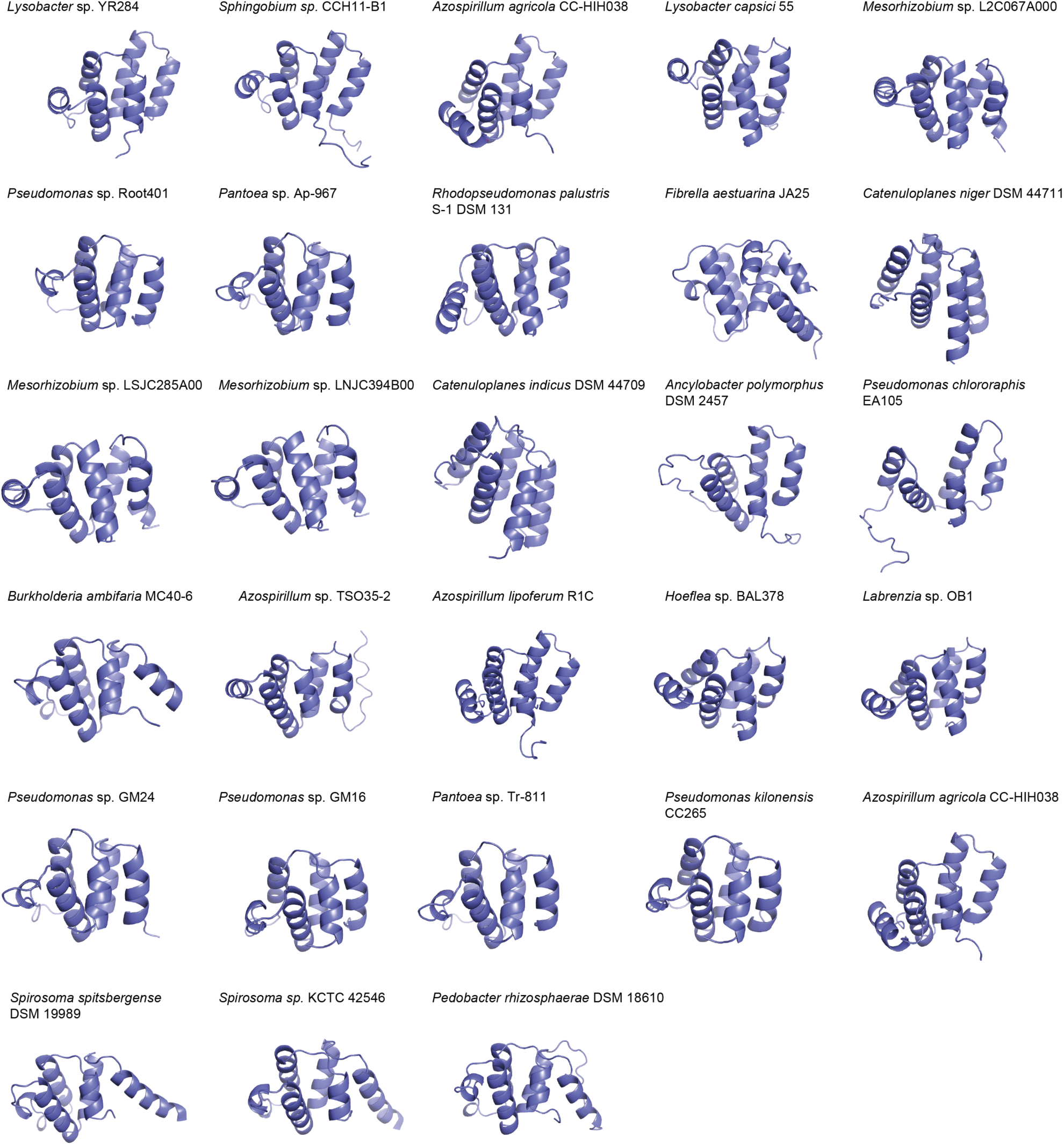
Predicted structures of CARD-like domains. AlphaFold2 prediction of N-terminal CARD-like domains from the proteases homologous to the protease of the *Lysobacter* gasdermin system (Table S2).

**Figure S5.**
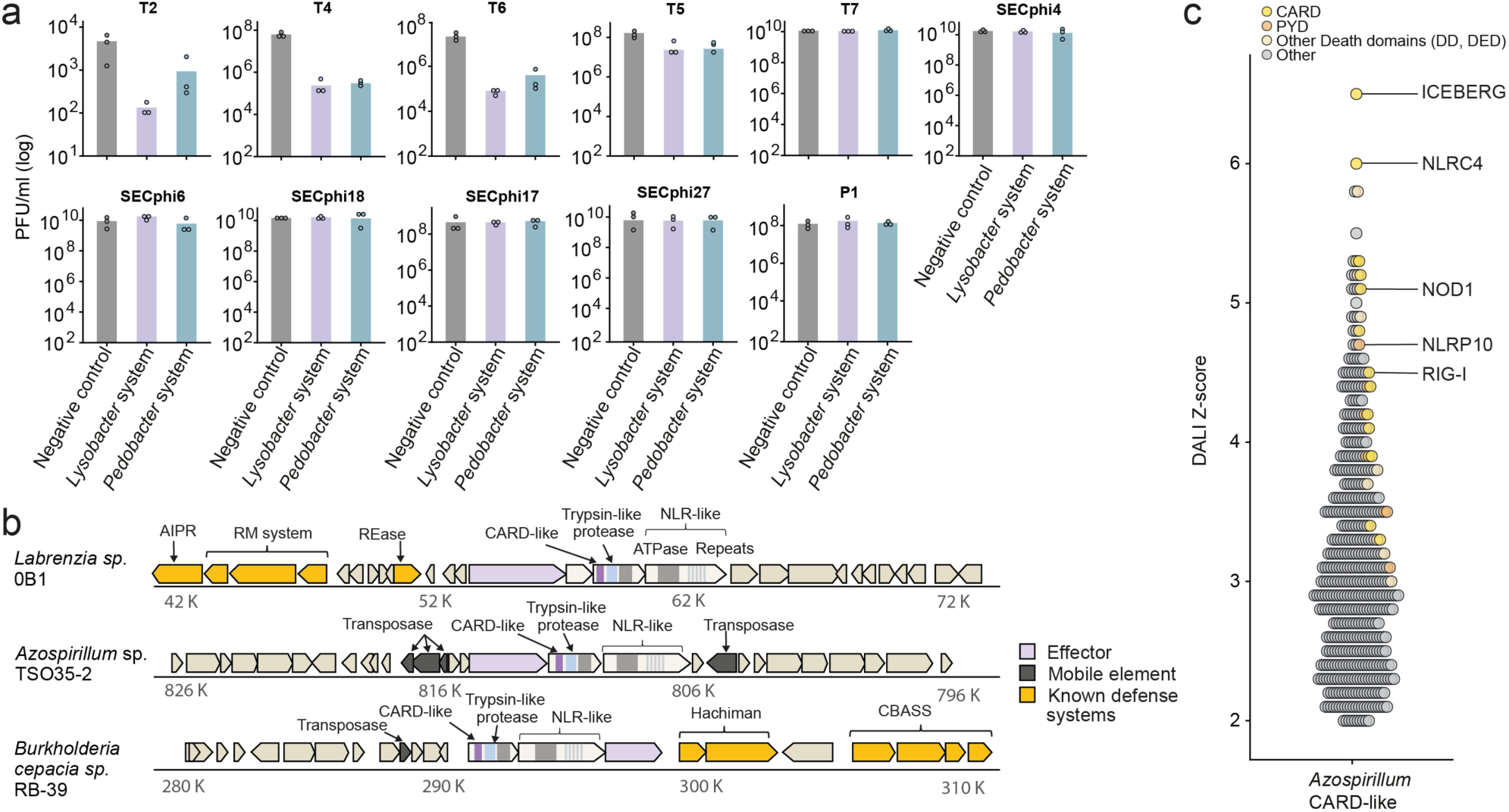
Systems homologous to the *Lysobacter* gasdermin system. **a.** Efficiency of plating of phages infecting *E. coli* cells that express the *Lysobacter* or *Pedobacter* defense systems. Negative control is a strain in which GFP is expressed instead of the gasdermin system. Data represent plaque-forming units (PFU) per milliliter. Average of three independent replicates, with individual data points overlaid. **b.** Representative instances of homologous systems in their genomic neighborhood. Genes known to be involved in anti-phage defense are shown in yellow (RM, restriction-modification; CBASS, cyclic-oligonucleotide-based antiphage signaling system; AIPR, abortive infection phage resistance; REase, restriction endonuclease). **c.** DALI Z score of protein structures similar to the structure of the *Azospirillum* CARD-like structure.

**Figure S6.**
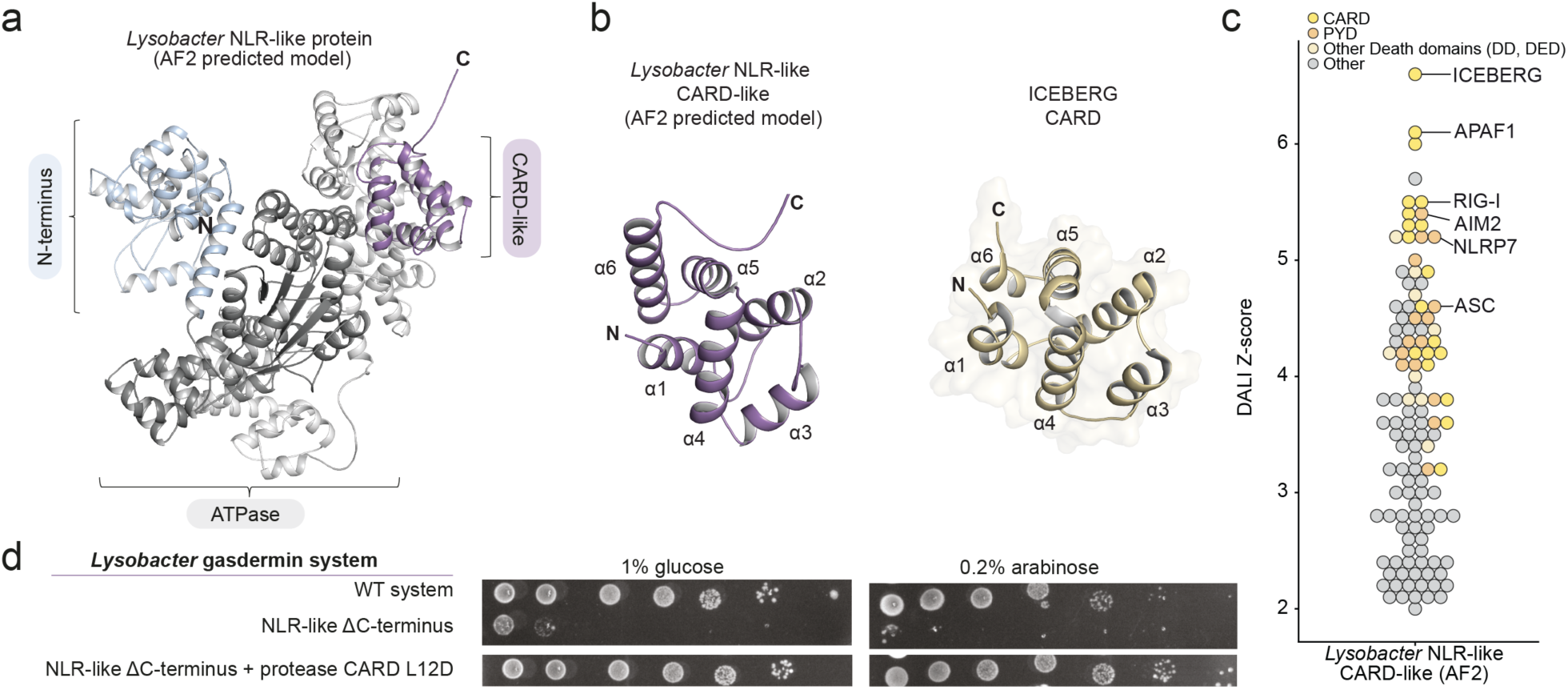
*Lysobacter* NLR-like protein encodes a CARD-like domain at the C-terminus. **a.** AlphaFold2 model of the NLR-like protein from the *Lysobacter* gasdermin system. **b.** AlphaFold2 model of the C-terminal domain of the NLR-like protein compared to the human ICEBERG CARD domain (PDB ID 1DGN). **c.** DALI Z scores of protein structures similar to the AlphaFold2 model of the CARD-like domain from the C-terminus of the *Lysobacter* NLR-like protein. **d.** Deletion of the C-terminus of the NLR-like gene in the *Lysobacter* gasdermin system leads to toxicity. Bacteria expressing the WT or mutated *Lysobacter* gasdermin system were plated in 10-fold serial dilution on LB-agar plates in conditions that repress expression (1% glucose) or induce expression (0.2% arabinose).

**Figure S7.**
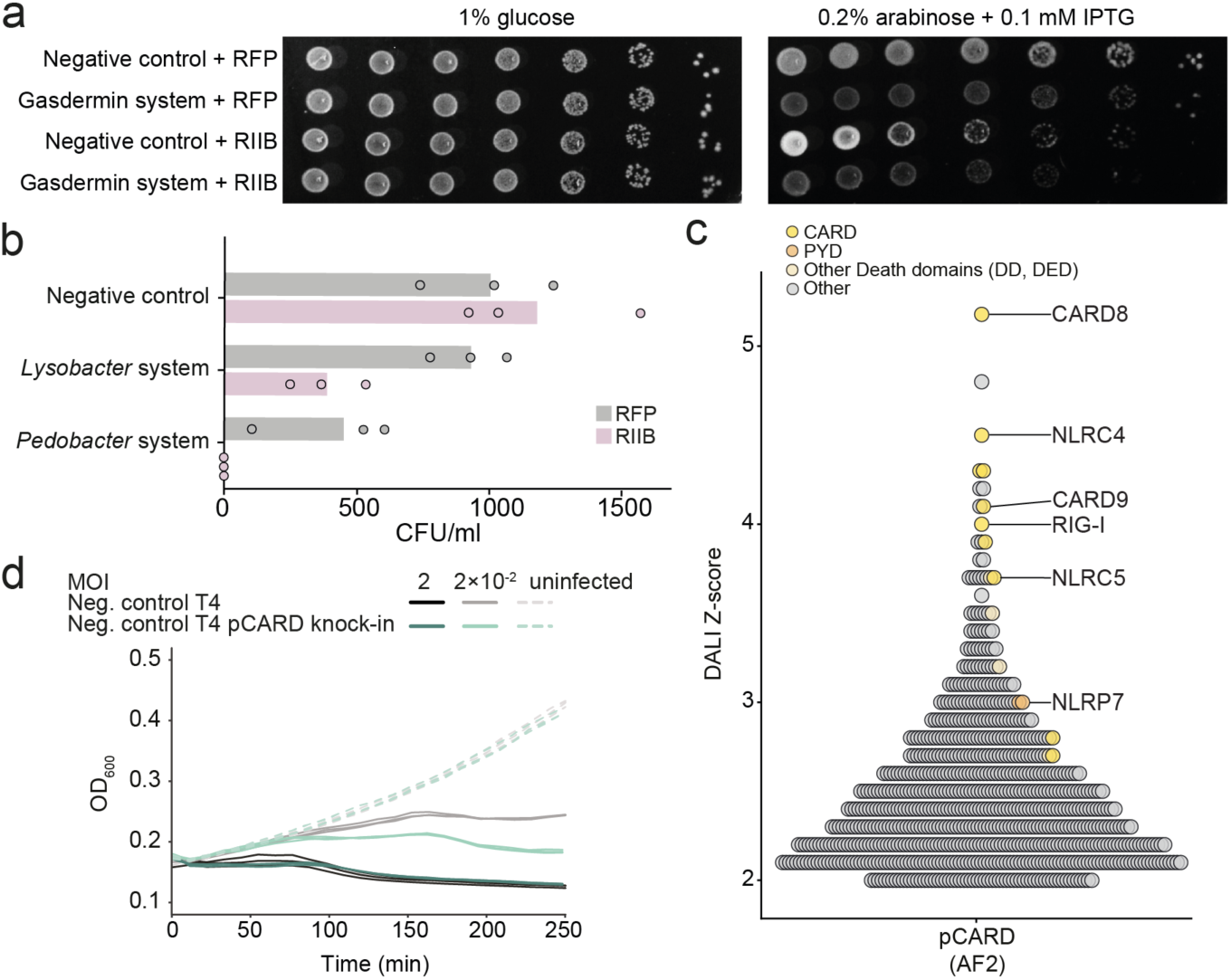
Phage proteins that activate or repress gasdermin-mediated defense. **a.** Bacteria expressing RIIB or RIIB together with the *Lysobacter* gasdermin system were plated in 10-fold serial dilution on LB-agar plates in conditions that repress expression (1% glucose) or induce expression (0.2% arabinose and 0.1 mM IPTG). **b.** Transformation efficiency assays of plasmids encoding *rIIB* or RFP into cells that contain the *Lysobacter* or *Pedobacter* systems on a pBAD plasmid. **c.** DALI Z scores of protein structures similar to the AlphaFold2 model of the phage CARD-only protein (pCARD). **d.** Liquid culture growth of *E. coli* control cells infected with WT T4 or with T4 engineered to encode the phage CARD-only gene (pCARD) instead of its *IPI* gene. Control bacteria, not encoding the defense system, were infected at time 0 at an MOI of 2 or 0.02 at 25 °C. Three independent replicates are shown for each MOI, and each curve represents an individual replicate.

### Supplementary Tables

**Table S1**. Crystallographic statistics.

**Table S2**. Systems that are homologous to the gasdermin-associated protease and ATPase proteins.

**Table S3**. Systems with structural homology to the gasdermin-associated protease.

### Supplementary Movies

**Movie S1**. Time-lapse microscopy corresponding to Figure 1a of live *E. coli* cells expressing GFP (green) mixed with cells expressing the *Lysobacter* gasdermin system (black). Cells were infected with phage T6 in the presence of propidium iodide (PI) and incubated at room temperature on an agar pad. Overlay images of phase contrast, green channel (GFP), and magenta channel (PI) are presented. Frames were taken every 3 minutes.

**Movie S2**. Time-lapse microscopy corresponding to Figure 1d of live *E. coli* cells expressing the WT *Lysobacter* gasdermin system in which gasdermin was N-terminally fused to GFP. Cells were infected with phage T6 in the presence of propidium iodide (PI) and incubated at room temperature on an agar pad. Overlay images of phase contrast, green channel (GFP), and magenta channel (PI) are presented. Frames were taken every 3 minutes.

**Movie S3**. Time-lapse microscopy corresponding to Figure 4c of live *E. coli* cells co-expressing RIIB with the WT *Lysobacter* gasdermin system, in which gasdermin was N-terminally fused to GFP. Cells were visualized at room temperature on an agar pad in the presence of propidium iodide (PI). Overlay images of phase contrast, green channel (GFP), and magenta channel (PI) are presented. Frames were taken every 3 minutes.

## Methods

### Bacterial strains, phages and growth conditions

The strain *Escherichia coli* NEB 5-alpha (New England Biolabs, NEB C2987H) was used as a cloning strain. *E. coli* K-12 MG1655 (ATCC 47076) was used in all experiments involving phage or toxicity. Unless mentioned otherwise, cells were grown in MMB (LB + 0.1 mM MnCl_2_ + 5 mM MgCl_2_, with or without 1.5% agar) with the appropriate antibiotics: ampicillin (100 µg/ml) or chloramphenicol (30 µg/ ml). Expression was repressed using 1% glucose and induced using 0.02%-0.2% arabinose or 0.1-1 mM IPTG.

Phage infections were performed in MMB media with or without 0.5% agar. *E. coli* phages P1, T4, T5, T7 and Lambda were provided by U. Qimron. Phages SECphi17, SECphi18, SECphi27, SECphi4, and SECphi6 were isolated in the Sorek laboratory^47, 48^. Phages T2 and T6 were obtained from the Deutsche Sammlung von Mikroorganismen und Zellkulturen (DSMZ) (DSM 16352 and DSM 4622, respectively).

### Plasmid construction

The *Lysobacter enzymogenes* gasdermin and the *Pedobacter rhizosphaerae* system were both expressed under an arabinose-inducible P_BAD_ promoter in the pBad/His A plasmid (Thermo Scientific V43001). The *Lysobacter* gasdermin system was previously cloned^10^ and the *Pedobacter rhizosphaerae* system was amplified from genomic DNA ordered from the DSMZ (DSM 18610, IMG gene accessions in Table S2) and cloned using the NEBuilder HiFi DNA Assembly cloning kit (NEB E5520S). As a negative control, the same plasmid encoding GFP was used^49^, unless stated otherwise.

The *rIIB* gene was amplified from phage T6 DNA (NCBI protein accession YP_010067419, nucleotide position: NC_054907, 167748-168686) and *rexAB* from phage Lambda DNA (NCBI accession AAA96580 and AAA96579, nucleotide position: J02459, 35825-37114). Both *rIIB and rexAB* were cloned under the IPTG-inducible P_LlacO-1_ promoter into the vector pBbA6c-RFP^50^ (Addgene 35290) using NEBuilder HiFi DNA Assembly. The gene encoding the phage CARD-only protein (pCARD) from *Acinetobacter* phage 133 (IMG gene accession 651703305) was synthesized and cloned by Genscript Corp. into the pBbA6c-RFP plasmid. As negative control, the original pBbA6c-RFP^50^ plasmid encoding an RFP gene was used.

GFP was fused to the N-terminus of gasdermin in the *Lysobacter* gasdermin system using NEBuilder HiFi DNA Assembly. The HA protein tag and point mutations were introduced using the KLD Enzyme mix (NEB M0554S) and were verified by Sanger sequencing and whole-plasmid sequencing (Plasmidsaurus).

### Phage infections assays

Phages were amplified from a single phage plaque using a liquid culture of *E. coli* MG1655 grown at 37°C to an OD_600_ of 0.3 in MMB media, incubated with the phage until culture collapse. The collapsed culture was centrifuged for 10 min at 3,200×g and the supernatant was filtered through a 0.2 µm filter.

Plaque assays were performed using the small-drop plaque assay method as previously described^47, 51^. Briefly, to measure defense, bacteria (*E. coli* MG1655 with system) and negative control (*E. coli* MG1655 with pBAD-GFP) were grown overnight at 37°C, diluted 1:100 and grown to an OD_600_ 0.5. 300 µl of the bacterial culture were mixed with 30 ml melted MMB agar (LB supplemented with 0.1 mM MnCl_2_, 5 mM MgCl_2_, 0.5% agar and 0.02% or 0.2% arabinose) and left to dry for 1 h at room temperature. 10-fold serial dilutions in MMB were performed for each of the tested phages and 10 µl was dropped on the bacterial layer. Plates were incubated overnight at 25°C. Plaque forming units (PFUs) were determined by counting the derived plaques after overnight incubation. When individual plaques could not be deciphered, a faint lysis zone across the drop area was considered to be ten plaques.

### Growth assays

*E. coli* MG1655 cells were grown at 37°C overnight in LB media supplemented with antibiotics and 1% glucose to avoid leaky expression. For the colony forming unit (CFU) count, 10-fold serial dilutions in 1×phosphate-buffered saline (PBS) were performed for each of the samples and 5 µl drops were put on LB-agar plates containing the appropriate antibiotics as well as 1% glucose and on LB-agar plates containing either 0.2% arabinose 1 mM IPTG, or both 0.2% arabinose and 0.1 mM IPTG. Plates were incubated overnight at 37°C and the bacterial colonies were imaged.

### Liquid infection assays

Overnight cultures of *E. coli* MG1655 encoding the WT or mutated *Lysobacter* gasdermin systems or control plasmids were diluted 1:100 in MMB with ampicillin and 0.2% arabinose and grown to an OD_600_ of 0.3. Afterwards, 180 µl of the cultures was transferred to a 96-well plate containing 20 µl of either 1× PBS (for uninfected samples) or phage lysate of T6 or T4 at various multiplicities of infection (MOIs). Plates were incubated at 25°C with shaking in a Tecan Infinite200 plate reader and the OD_600_ was measured every 10 min.

Propagation of phage T6 on control and gasdermin-expressing cells was measured by infecting exponential cultures at OD_600_ ∼0.3 with T6 at an MOI of ∼0.01. The infected cultures were incubated at 25°C with shaking and sampled at the indicated time points, centrifuged at 14,000×g for 1 min and the free phage titer was assessed in the supernatant by plaque assays on *E. coli* MG1655, as described above.

### Isolation of mutant phages that overcome defense

To isolate mutant phages that escape gasdermin-mediated defense, phages were plated on bacteria expressing the gasdermin system using the double-layer plaque assay^52^. Bacterial cells were grown in MMB (supplemented with 0.2% arabinose) to an OD_600_ of 0.3. Then, 100 µl of bacterial cells was mixed with 100 µl phage lysate and left at room temperature for 10 min. Afterwards, 5 ml of pre-melted 0.5% MMB agar (supplemented with 0.2% arabinose) was added and the mixture was poured onto a bottom layer of 1.5% MMB agar. The double-layer plates were incubated overnight at 25°C and single plaques were picked into 90 µl of 1×PBS. Phages were left for 1 h at room temperature during which the phages were mixed several times by vortex to release them from the agar into the phage buffer. To test the ability of the picked phages to escape from the defense provided by the *Lysobacter* gasdermin system, the small-drop plaque assay was used. For this, 10-fold serial dilutions of the WT phage and the phage plaques picked from the gasdermin-expressing strain were plated on bacteria harboring the gasdermin system, the *Pedobacter* system or a negative control expressing GFP (as described above^51^). PFUs were counted after overnight incubation at 25°C. Isolated phages for which there was decreased defense compared with the ancestor phage were further propagated by picking a single plaque formed on the defense gene in the small-drop plaque assay into 2 ml liquid culture of *E. coli* at an OD_600_ of 0.3. The phages were incubated with the bacteria at 37°C for 3 h. The lysates were then centrifuged at 14,000×g for 10 min, and the supernatant was filtered through a 0.2 µm filter to get rid of the remaining bacteria. The phage titer was checked using the small-drop plaque assay on the negative-control strain as described above.

### Sequencing and genome analysis of phage mutants

High-titer phage lysates (>10^7^ PFU/ml) of the original T6 phage and the isolated phages were used for DNA extraction. 0.5 ml of phage lysate was treated with DNase-I (Merck catalogue no. 11284932001) added to a final concentration of 20 µg/ml and incubated at 37 °C for 1 h to remove bacterial DNA. DNA was extracted using the QIAGEN DNeasy blood and tissue kit (catalogue no. 69504). Libraries were prepared for Illumina sequencing using a modified Nextera protocol as previously described^53^. Following sequencing on Illumina NextSeq500, reads were aligned to the phage reference genome (NCBI accession: NC_054907.1) and mutations compared with the reference genome were identified using Breseq (v.0.29.0 or v.0.34.1) with default parameters^54^. Only mutations that occurred in the isolate mutants, but not in the ancestor phage, were considered.

### Transformation efficiency

Strains of *E. coli* carrying either the *Pedobacter* system, the *Lysobacter* gasdermin system or GFP as a negative control were used as recipients for chemical transformation in TSS media^55^. An overnight culture of the recipient bacteria grown in MMB was diluted 1:100 into 5 ml of fresh media and grown at 37°C to an OD_600_ of 0.2. Bacteria were then centrifuged at 4000×g for 5 min, after which the bacterial pellet was resuspended on ice in 100 µl cold TSS media (LB supplemented with 10% (w/v) PEG 8000, 0.6% (w/v) MgCl_2•_6H_2_0, and 5% (v/v) DMSO). 50 µl of the bacteria were transferred into a fresh tube and 100 ng of the vector expressing the phage *rIIB* gene or a negative control vector expressing RFP was added. The bacteria were incubated with the vector on ice for 5 min, then at room temperature for 5 min, and again on ice for 5 min. 950 µl MMB was added and the bacteria were incubated at 37°C for 1 h. 10-fold serial dilutions of the bacteria in PBS, were then plated on LB-agar plates supplemented with 100 µg/ml ampicillin, 30 µg/ml chloramphenicol and 1% glucose. Transformation efficiency was assessed by counting single colonies formed after overnight incubation at 37°C.

### Bacterial live cell imaging

*E. coli* strains were grown overnight in MMB media supplemented with the appropriate antibiotics at 37°C. The overnight cultures were diluted 1:100 into fresh media and incubated shaking at 25°C until the bacterial cultures reached an OD_600_ of 0.4. Then, 300 µl of bacteria was either infected with phage T6 (MOI=2) and incubated for 10 min at room temperature or directly washed by centrifugation at 5,000×g for 5 min and resuspension in 5 µl of 1×PBS supplemented with 0.2 µg/ml propidium iodide (PI) (Sigma-Aldrich P4170). Cells were placed on a 1% agarose pad that was supplemented with either 0.2% arabinose or both 0.2% arabinose and 0.1 mM IPTG. Cell images were acquired over time every 3 minutes using an Axioplan2 microscope (Zeiss) equipped with ORCA Flash 4.0 camera (Hamamatsu). Microscope control and image processing were carried out with Zen software version 2.0 (Zeiss).

### Generation of phage knock-in using Cas13a

The gene encoding the phage CARD-only protein (pCARD) was introduced into the phage T4 genome using a CRISPR-based selection strategy as described previously^37^. Briefly, a plasmid serving as the template for homologous recombination of pCARD into the T4 genome was created by cloning the pCARD gene between 50 bp recombination overhang sequences (ordered from Twist Biosciences) in the pGEM-9Zf plasmid (Promega P2391). The recombination overhangs targeted the pCARD gene for insertion after the predicted promoter of the T4 *IPI* gene^38^, replacing the *IPI* gene in the mutated phage. The plasmid pBA559 (Addgene 186235), encoding the LbuCas13a gene under the TetR inducible promoter^37^, was used to clone the guide RNA (5′-cagaagtaagagtagcttcggtaatggtag-3′) targeting the T4 wildtype *IPI* gene. The guide was cloned into pBA559 using BsaI (NEB R3733L) restriction cloning. Both plasmids were introduced into *E. coli* MG1655. Cells carrying the plasmid encoding for the recombination template were grown until an OD_600_ of 0.3. The cells were infected with T4 at an MOI of 0.01 and incubated at 37°C with constant shaking. After culture collapse (around 3 h post infection) the culture was centrifuged at 3,200×g for 5 min and the supernatant was filtered through a 0.2 µm filter. The filtered phage was then used to infect cells carrying the pBA559 *IPI* guide RNA plasmid in a small-drop plaque assay (described above). Single plaques formed after overnight incubation at 37°C were picked into 100 µl of 1×PBS and checked for the phage pCARD knock-in using Sanger sequencing. Positive phage T4 clones encoding for pCARD went through three rounds of plaque purification before generating a high-titer stock. The T4 pCARD knock-in phage was then used to infect cells expressing the *Lysobacter* gasdermin system in liquid infection assays (described above).

### Protein gel electrophoresis and Western blotting

Cell cultures expressing the HA-tagged *Lysobacter* gasdermin system, HA-tagged mutated *Lysobacter* gasdermin system or the WT *Lysobacter* gasdermin system were grown at 25°C in MMB with 0.2% arabinose until they reached an OD_600_ of 0.5. 1 ml of the cultures was sampled, centrifuged at 3,200×g for 5 min and the supernatant was discarded. The remaining cultures were infected with phage T6 at an MOI of 2. The cultures were sampled again 80 min after infection, centrifuged, and the supernatant was discarded. Cells expressing both the HA-tagged *Lysobacter* gasdermin system and the phage RIIB protein were grown at 25°C in MMB with 1% glucose until an OD_600_ of 0.5. Then, 1 ml of the cultures was sampled, centrifuged, and the supernatant was removed. Expression of the gasdermin system and RIIB was subsequently induced with 0.2% arabinose and 1 mM IPTG. Cultures were sampled again 80 min after induction.

Lysates were resuspended in 1×Bolt LDS Sample Buffer (Thermo Scientific B0007) supplemented with 50 mM DTT. The samples were boiled at 95°C for 5 min and 10 µl of each sample was separated by 4-12% Bis-Tris SDS-PAGE (Thermo Scientific NW04122BOX) in 1×MES buffer (Thermo Scientific B0002) for 22 min at 200 V. For Western-blotting, the gel was transferred to a nitrocellulose membrane (Invitrogen LC2001) for 1 h at 10 V in 1×transfer buffer (Thermo Scientific BT00061) and probed with a primary mouse anti-HA antibody (Sigma Aldrich H9658; 1:3000 dilution in TBS-T with 3% BSA). Visualization of the primary antibody was performed using HRP-conjugated goat anti-mouse secondary antibody (Thermo Scientific 31430; 1:10,000 dilution in TBS-T) and incubation with ECL solution (Merck Millipore WBLUF0500).

### Recombinant protein expression and purification

DNA encoding putative CARD-like domains was synthesized as codon-optimized gene fragments (IDT) and cloned by Gibson assembly into a custom pET plasmid fused to a N-terminal 6×His-hSUMO2 tag. Plasmids were verified by Sanger sequencing, transformed into BL21 CodonPlus(DE3)-RIL *E. coli*, selected on antibiotic-containing LB agar, and colonies were used to inoculate starter cultures in MDG media (0.5% glucose, 25 mM Na_2_HPO_4_, 25 mM KH_2_PO_4_, 50 mM NH_4_Cl, 5 mM Na_2_SO_4_, 2 mM MgSO_4_, 0.25% aspartic acid, 100 mg/ml ampicillin, 34 mg/ml chloramphenicol, and trace metals) for overnight growth at 37°C. The MDG starter cultures were used to inoculate M9ZB media (0.5% glycerol, 1% Cas-amino Acids, 47.8 mM Na_2_HPO_4_, 22 mM KH_2_PO_4_, 18.7 mM NH_4_Cl, 85.6 mM NaCl, 2 mM MgSO_4_, 100 mg/ml ampicillin, 34 mg/ml chloramphenicol, and trace metals) and cultures were grown at 37°C with 230 RPM shaking to an OD_600_ of ∼2.5. Cultures were cooled on ice for 20 min and then expression was induced by the addition of 0.5 mM IPTG before incubation overnight at 16°C with shaking at 230 RPM. For each putative CARD-like domain, 1-2× 1 liter cultures were expressed, pelleted, flash frozen in liquid nitrogen, and stored at −80°C.

Protein purification was performed at 4°C with buffers containing 20 mM HEPES-KOH (pH 7.5) and salts as indicated. Frozen cell pellets from protein expression were thawed on ice and lysed by sonication in a buffer containing 400 mM NaCl and 1 mM DTT. The lysate was clarified by centrifugation and passing through glass wool, bound to NiNTA agarose resin (QIAGEN), washed with buffer containing 1 M NaCl and 1 mM DTT, and eluted with buffer containing 400 mM NaCl, 300 mM imidazole (pH 7.5), and 1 mM DTT. Recombinant hSENP2 protease (D364–L589, M497A) was added to the NiNTA elution to cleave off the SUMO2 tag, and the eluate was dialyzed overnight in buffer containing 125–250 mM KCl and 1 mM DTT. The dialyzed protein was subsequently purified by size-exclusion chromatography with a 16/600 Superdex 75 column (Cytiva) pre-equilibrated with a buffer containing 250 mM KCl and 1 mM TCEP. Sized proteins were concentrated to >50 mg/ml using 10 kDa MWCO concentrators (Millipore), flash frozen on liquid nitrogen, and stored at −80°C. SeMet-substituted proteins were prepared in M9 media and purified in buffers containing 1 mM TCEP in place of DTT.

### Crystallization and structure determination

Putative CARD-like domains from proteases and NLR-like proteins were screened for optimal expression and solubility. None of the 15 CARD-like domains screened from NLR-like proteins yielded soluble proteins for use in crystallography (Table S2). In contrast, most CARD-like domains from the proteases were found to express well and be soluble, and the *Lysobacter enzymogenes* (IMG gene ID 2841794909) and *Azospirillum sp.* (IMG gene ID 2882854644) CARD-like domains were used for crystallization. Crystals were grown by the hanging-drop vapor diffusion method at 18°C with NeXtal crystallization screens and optimized with EasyXtal 15-well trays (NeXtal). Proteins for crystallization were thawed from −80°C stocks on ice and diluted to concentrations of 20 mg/ml protein and a final buffer of 20 mM HEPES-KOH (pH 7.5), 60 mM KCl, and 1 mM TCEP. 15-well crystal trays were set with 2 µl drops containing diluted protein and reservoir at a 1:1 ratio over wells with 350 µl reservoir solution. Crystals were grown for 2-3 days, incubated with select cryoprotectant, and harvested by flash freezing crystal harvest loops in liquid nitrogen. The CARD-like domain from the *Lysobacter* protease (amino acids 2–92) crystals grew in 3.3 M sodium formate and were cryoprotected in reservoir solution. The SeMet-substituted *Lysobacter* CARD-like domain crystals grew in 3.3 M sodium formate and were cryoprotected in 5 M sodium formate. The CARD-like domain from the *Azospirillum* trypsin-like protein (amino acids 13–111) crystals grew in 100 mM HEPES-KOH (pH 7.5), 200 mM ammonium acetate, and 25% PEG-3350 and were cryoprotected in the same solution supplemented with 20% ethylene glycol. X-ray diffraction data were acquired using Northeastern Collaborative Access Team beamlines 24-ID-C and 24-ID-E (P30 GM124165), and used a Pilatus detector (S10RR029205), an Eiger detector (S10OD021527) and the Argonne National Laboratory Advanced Photon Source (DE-AC02–06CH11357).

Data were processed with XDS and AIMLESS using the SSRL autoxds script (A. Gonzalez)^56^. The *Lysobacter* CARD-like structure was phased with anomalous data from SeMet-substituted crystals using Phenix Autosol version 1.19^57, 58^. The *Azospirillum* CARD-like structure was phased by molecular replacement using an AlphaFold2 model and Phaser-MR in Phenix version 1.19^57, 58^. Atomic models were built in Coot^59^ and refined in Phenix. Statistics were analyzed as described in Table S1^60–62^. Structure data were deposited in the Protein Data Bank (PDB IDs 8SRZ and 8SS1). Structure figures were generated using PyMOL version 2.4.0 (Schrödinger, LLC).

### Identification of homologous systems and sequence alignment

The *Lysobacter* protease and ATPase protein sequences (Table S2) were used to search for homologs using BLASTp against the IMG database^46^ with default parameters (Table S2). Sequence alignments were performed using MAFFT v.7.402^63^ with default parameters and structure-guided alignments were performed using PROMALS3D^64^. The alignments were visualized using Jalview^65^.

### Structural predictions

AlphaFold2^15^ was used to model the structure of homologous proteases (Table S2, S3). To locate the CARD domain coordinates within protease homologs the SWORD (v1.0) tool^66^ was run on the AlphaFold2 structural models. The derived domains were manually inspected for the presence of the six characteristic alpha-helices (Figure S4). For domain annotation, the predicted structures were further analyzed using the DALI webserver^20^. Predicted structure figures were generated using PyMOL version 2.4.0 (Schrödinger, LLC).

### Structural search for CARD-encoding defense systems

The AlphaFold2 model of the *Lysobacter* protease was used as a query to search the AlphaFold/ UniProt50 database using the Foldseek webserver^24^ in TM-align mode. UniProt50 hits with probability >0.4 were used to find protein sequence homologs using MMseqs2^67^ (release 6-f5a1c) with default parameters, by searching them against the cluster representatives of a database of proteins from 38,167 bacterial and archaeal genomes that was previously downloaded^68^ from the Integrated Microbial Genomes (IMG) database. For each UniProt50 sequence, the protein cluster of the best hit was examined manually and further analyzed for the presence of conserved gene cassettes (Table S3).

## Data Availability

Data that support the findings of this study are available within the article and its Supplementary Tables. IMG accessions appear in Supplementary Tables 2 and 3. Coordinates and structure factors of the *Lysobacter* and *Azospirillum* CARD-like domains have been deposited in the PDB under the accession codes 8SRZ and 8SS1, respectively.

## Acknowledgements

The authors thank the Sorek and the Kranzusch laboratory members for comments on the manuscript and fruitful discussion. R.S. was supported, in part, by the European Research Council (grant no. ERC-AdG GA 101018520), Israel Science Foundation (MAPATS Grant 2720/22), the Deutsche Forschungsgemeinschaft (SPP 2330, Grant 464312965), the Ernest and Bonnie Beutler Research Program of Excellence in Genomic Medicine, and the Knell Family Center for Microbiology. P.J.K. was supported, in part, by the Pew Biomedical Scholars program, The Mathers Foundation, the Parker Institute for Cancer Immunotherapy, and the National Institutes of Health (1DP2GM146250). T.W. was supported by a Minerva Foundation postdoctoral fellowship and by a European Molecular Biology Organization (EMBO) postdoctoral fellowship (ALTF 946-2020). A.G.J was supported by a Life Science Research Foundation postdoctoral fellowship of the Open Philanthropy Project. A.M. was supported by a fellowship from the Ariane de Rothschild Women Doctoral Program and, in part, by the Israeli Council for Higher Education via the Weizmann Data Science Research Center. E.Y. was supported, in part, by the Israeli Council for Higher Education (CHE) via the Weizmann Data Science Research Center. X-ray data were collected at Northeastern Collaborative Access Team beamlines 24-ID-C and 24-ID-E (no. P30 GM124165), including use of a Pilatus detector (no. S10RR029205), an Eiger detector (no. S10OD021527) and the Argonne National Laboratory Advanced Photon Source (no. DE-AC02-06CH11357).

## Author Contribution

The study was designed by T.W., A.G.J., P.J.K. and R.S. T.W. performed all genomic analyses, cellular imaging, phage infection experiments and gasdermin cleavage assays. A.G.J. performed all X-ray crystallography and structural analyses. A.M., E.Y. and K.L. performed genomic analyses of bacterial defense systems and phage CARD-like proteins. R.H., J.G., and T.W. engineered the pCARD protein into phage T4. F.S., A.B.H., and A.G.J screened candidate CARD-like protein expression *in vitro*. T.W., A.G.J., P.J.K., and R.S. wrote the manuscript. R.S. and P.J.K. supervised the study.

