## Supplementary material for "CARD-like domains mediate anti-phage defense in bacterial gasdermin systems": Table S1

**Table S3. Crystallographic statistics, Related to Figure 2 and Figure 3**

|  | ***Lysobacter* bCARD** | ***Lysobacter* bCARD**  (SeMet) | ***Azospirillum* bCARD** |
| --- | --- | --- | --- |
| **Data Collection** | | |  |
| Resolution (Å)^a^ | 46.64–1.25 (1.25–1.27) | 46.66–1.45 (1.45–1.47) | 36.24–1.88 (1.88–1.92) |
| Wavelength (Å) | 0.9792 | 0.9792 | 0.9792 |
| Space group | P 6_5_ | P 6_5_ | P 1 2_1_ 1 |
| Unit cell: a, b, c (Å) | 58.25 58.25 122.35 | 58.31, 58.31, 122.14 | 37.96, 65.37, 38.83 |
| Unit cell: α, β, γ (°) | 90.00 90.00 120.00 | 90.00, 90.00, 120.00 | 90.00, 111.06, 90.00 |
| Molecules per ASU | 2 | 2 | 2 |
| Total reflections | 1,106,507 | 867,147 | 91,309 |
| Unique reflections | 64,141 | 41,615 | 14,427 |
| Completeness (%)^a^ | 99.0 (96.1) | 100.0 (99.3) | 99.1 (91.4) |
| Multiplicity^a^ | 17.3 (16.5) | 20.8 (19.4) | 6.3 (6.0) |
| *I/σI^a^* | 23.0 (2.1) | 10.7 (1.3) | 7.7 (1.4) |
| CC(1/2)^b^ (%)^a^ | 100.0 (88.1) | 99.8 (67.7) | 99.6 (52.6) |
| Rpim^c^ (%)^a^ | 1.4 (27.1) | 4.0 (74.2) | 6.2 (79.3) |
| Sites |  | 1 |  |
| **Refinement** | | |  |
| Resolution (Å) | 39.92–1.25 |  | 36.24–1.88 |
| Free reflections | 64,038 |  | 14,258 |
| R-factor / R-free | 15.9 / 17.8 |  | 20.4 / 24.1 |
| Bond distance (RMS Å) | 0.015 |  | 0.004 |
| Bond angles (RMS °) | 1.257 |  | 0.612 |
| **Structure/Stereochemistry** | | |  |
| No. atoms: protein | 1,338 |  | 1,535 |
| No. atoms: water | 325 |  | 121 |
| Average B-factor: protein | 22.4 |  | 29.3 |
| Average B-factor: water | 33.6 |  | 38.4 |
| Ramachandran plot: favored | 100.00% |  | 100.00% |
| Ramachandran plot: allowed | 0.00% |  | 0.00% |
| Ramachandran plot: outliers | 0.00% |  | 0.00% |
| Rotamer outliers | 0.00% |  | 1.91 |
| MolProbity^d^ score | 1.36 |  | 1.39 |
| Protein Data Bank ID | 8SRZ |  | 8SS1 |

^a^ Highest resolution shell values in parentheses

^b^ (Karplus and Diederichs, 2012)

^c^ (Weiss, 2001)

^d^ (Chen et al., 2010)
